# Maternal singing synchronizes the preterm infants’ brain

**DOI:** 10.1101/2025.06.24.660986

**Authors:** D. Benis, J. Margaria, F. Barcos-Munoz, J. Miehlbradt, L. Lavezzo, D. Grandjean, A. Vollenweider, S. Henriot, P. Hüppi, M. Filippa

## Abstract

Singing to infants is a universal human practice that has beneficial effects on infant’s cognitive and affective development. Children born preterm have impaired brain development, and their perception of maternal speech is known to be affected by the atypical hospital auditory environment. Understanding how preterm infants perceive maternal singing is of critical importance, yet it remains largely unexplored. Using high-density EEG, we examined neural responses to the same melody presented through maternal singing, stranger singing, and instrument, and compared the responses in 12 preterm infants. Moreover, to examine their processing of spatialisation, auditory stimuli were presented under monaural and binaural conditions.

Preterm newborns are able to discriminate the same melody when sung by their mother, a stranger, or played by an instrument. When presented monaurally, the mother’s singing voice enhances widespread brain synchrony across the entire scalp. In contrast, this synchrony diminishes with binaural spatialization. These findings suggest that maternal singing constitutes a highly salient auditory stimulus for preterm newborns, eliciting a distinct neural signature. Given that brain synchrony is a critical component of healthy brain function and development, harnessing maternal singing may offer a promising, natural intervention to support neurodevelopment—particularly in vulnerable populations such as preterm infants.

## Introduction

Identifying and linking with caregivers is vital for organisms since the beginning of life, and human newborns largely rely on their auditory system for establishing these connections. Preterm infants’ brains develop in an atypical auditory environment, where maternal sounds are no longer the predominant stimuli they are exposed to. While differences in how preterm newborns process their mother’s speech compared to term infants have been studied (Adam-Darque et al., 2020; Filippa et al., 2023), their perception of maternal singing has yet to be explored. Whether maternal singing remains a salient and preserved stimulus for preterm infants is the central question of this investigation.

### Born too soon

Prematurity is recognized as a major clinical risk factor for neurodevelopmental disorders, leading to disrupted brain maturation (Cheong et al., 2021). Preterm infants are particularly vulnerable to brain lesions, and assessments conducted at term-corrected age reveal altered anatomical patterns (Dubois et al., 2008; Dubois et al., 2009; Hüppi et al., 1996). These include a range of macrostructural abnormalities, such as delayed differentiation of cerebral gray and white matter, impaired myelination, and white matter injuries (Ment et al., 2009). Such deviations from typical neurodevelopment observed in full-term infants often result in long-term cognitive and neurobehavioral challenges (Bell et al., 2022). These challenges are strongly associated with academic difficulties (Twilhaar et al., 2018) and an increased risk of psychiatric disorders, particularly among extremely preterm infants (Johnson et al., 2010; Treyvaud et al., 2013).

Both brain injury and impaired brain development—whether resulting from or occurring independently of injury—are significant contributors to adverse neurodevelopmental outcomes in preterm infants, and various factors may mediate these brain alterations associated with preterm birth (Inder et al., 2023). Among these factors, environmental sensory inputs, such as auditory and visual stimuli, play a critical role in shaping the trajectories of neuronal maturation (Pineda et al., 2014).

### Auditory environment and auditory development

The Neonatal Intensive Care Unit (NICU) auditory environment is characterized by elevated noise levels, predominantly from medical equipment, which can exceed recommended standards (Smith et al., 2018). Additionally, preterm infants in the NICU are exposed to fewer language interactions compared to full-term infants (Liszka et al., 2019), a fact which may impact auditory development and subsequent language acquisition (Vohr, 2014).

The auditory system develops in utero, with responses detectable as early as 19 weeks of gestation (Chang & Kanold, 2021). In response to auditory stimuli, the synchronous activation of inner hair cells is crucial for clustering primary auditory cortical neurons and establishing tonotopic sensory maps in the auditory cortex (Huberman et al., 2008). Extensive research highlights the role of peripheral sensory receptors in forming somatosensory maps, with activity-dependent plasticity refining these processes during the perinatal and early postnatal periods (Chang & Merzenich, 2003). Consequently, the prenatal experience with language and other intrauterine sounds positively shapes the brain (Mariani et al., 2023). Conversely, exposure to inappropriate sensory stimuli or noise in the NICU environment can interfere with brain maturation, hinder the development and fine-tuning of the auditory system, and lead to alterations in sensory cortical connectivity and function.

### Altered voice and sound perception in preterm newborns

The mature adult brain, has specialized regions for recognizing familiar voices (Maguinness et al., 2018), and decades of research have demonstrated that newborns can process their mother’s voice distinctively during pregnancy (Jardri et al., 2012) and shortly after birth (Beauchemin et al., 2011; DeCasper & Fifer, 1980).

Preterm infants equally possess minimal brain circuitry sufficient for attending to speech and processing its content. Early on, they can discriminate between syllables, reflecting an early specialization of the human cortex for speech processing. This specialization suggests that the neural architecture required for speech perception emerges early, prioritizing biologically significant stimuli such as the maternal voice (Mahmoudzadeh et al., 2013; Mahmoudzadeh et al., 2017).

However, at term-equivalent age, preterm infants exhibit atypical auditory processing compared to their full-term peers (Therien et al., 2004). When comparing terms versus preterm newborn’s speech perception (Adam-Darque et al., 2020), it has been found a similar activation in superior temporal gyrus, anterior cingulate cortex and orbitofrontal cortex for maternal and stranger voice in the two groups, but with very limited differences between mother and stranger voice perception for preterms. Full-terms, on the other hand, showed a larger contrast in the auditory paradigm between the two voices both in fMRI as well as in EEG (Adam-Darque et al., 2020).

Additionally, in preterm infants, the maternal voice triggered a significantly different neural response compared to the frequently heard stranger’s voice, occurring between 670 and 860 ms over the centro-parietal electrodes. This late positive mismatch response (MMR) has been interpreted as an early neural correlate of the adult P300b, indicating the emergence of voice differentiation mechanisms (Adam-Darque et al., 2020). Using the nurse’s voice as a comparison, Saito et al, demonstrated that in preterm newborns the mother’s and the nurse’s voices activated the left frontal area in both groups. However, only the nurse’s voice activated the right frontal area, highlighting not only that premature infants reacted differently to the two voice stimuli, but also that the nurses’ voices, to which preterm infants are exposed, could be linked to emotionally relevant or stressful experiences (Saito et al., 2009).

A general difference between preterms and full terms’ speech processing has been observed in the theta frequency band within the left temporal region, where preterms exhibited significant activation in response to strangers’ speech and term born infants for the mother’s speech (Filippa et al., 2023). Interestingly only full terms exhibited a late gamma band increase in response to maternal speech, potentially suggesting a more developed brain response, probably due to a more mature thalamocortical connectivity.

Distinct positive mismatch responses have also been observed in the two populations when exposed to speech and non-speech sounds, interpreted as auditory cortex immaturity in preterms (Kostilainen et al., 2020). Similarly, using mismatch negativity, researchers have identified atypical cortical auditory processing of non-vocal sounds in preterm infants (Fellman et al., 2004). For instance, Fellman et al. examined event-related potentials in preterm infants during their first year and found significantly reduced or absent mismatch negativity responses to deviant auditory stimuli (Fellman et al., 2004). This effect of prematurity is dependent on post-conceptual age at birth (deRegnier et al., 2002), with important impacts on speech prosody discrimination (Alexopoulos et al., 2021).

### Singing is universal and beneficial

Singing to infants is a culturally universal phenomenon known to regulate arousal (Cirelli et al., 2020; Nakata & Trehub, 2004; Trehub et al., 2015), alleviate distress (Corbeil et al., 2016), and regulate attention (Corbeil et al., 2013; Nguyen et al., 2023; Tsang et al., 2017), modulating infant’ sensorimotor, language and socioemotional development (Punamäki et al., 2024). It fosters social-communicative interpersonal engagement, by entraining audio-visual synchronization (Lense et al., 2022), thus potentially increasing information coupling between infant and adult brains (Leong et al., 2017). Despite variations in its acoustic forms and musical structures across cultures, it exhibits a set of universal acoustic features that distinguish it from both speech and non-infant-directed singing (Hilton et al., 2022). In the last decades, several early interventions demonstrated that the exposure to maternal speech and singing improves clinical outcomes, including physiological state, behavior, and neurological development (Filippa et al., 2017; Haslbeck et al., 2023; Provenzi et al., 2018). In particular, recurrent exposure to maternal singing during kangaroo care enhanced auditory processing (Partanen et al., 2022) and auditory discrimination of phonetic and emotional speech sounds (Kostilainen et al., 2021) in preterm infants at term age.

Preterm infants’ directed speech and singing elicit differential behavioural and physiological responses in preterms, with speech demonstrating stronger analgesic effects than singing, potentially due to a greater enhancement of oxytocin levels (Filippa et al., 2021). In contrast, singing has a more powerful impact on the maturation of the autonomic nervous system after repeated exposure during hospitalization (Filippa et al., 2024). It seems, therefore, that these two forms of communication with infants serve different functions in their ontogenesis and development (Corbeil et al., 2013; Trainor & Cirelli, 2015).

Speech and music perception in adults are supported by specific although partially overlapping frontal-medial-temporal and cerebellar network (Callan et al., 2007; LaCroix et al., 2015; Patel, 2003; Peretz et al., 2015). In adults, speech processing shares several common EEG event-related potential correlates with singing voice perception (Patel et al., 1998; Proverbio & Piotti, 2022). Interestingly, an iEEG study on epileptic patients showed that, while music and speech presented shared oscillatory power correlates in several brain structures, specificities were observed between the two modalities at the scale of frequency-specific functional networks, especially in the delta, theta and alpha bands (Te Rietmolen et al., 2024).

In young infants, only two studies to our knowledge have investigated the EEG correlates of singing voices. An oscillatory peak in the delta band as well as cortical tracking by cortical delta and alpha activity of nursery rhymes sung by non-caregiver adults were observed in 4 months old infants (Attaheri, Choisdealbha, et al., 2022; Attaheri, Panayiotou, et al., 2022). However, the oscillatory correlates of vocal singing in preterm neonates have not, to our knowledge, been investigated to date, nor has the impact of singing voice familiarity. This gap in the current literature is particularly significant, as singing represents a unique channel of communication that may be more exclusively shared between the infant and the primary caregiver, implicating a high degree of intimacy (Woodward, 2019). In the current study we intend to address this gap in literature by investigating in preterm infants the EEG oscillatory correlates of maternal singing compared to a stranger’s infant-directed singing and to a musical instrument.

### Relevance of neural synchrony

In the current study, we will focus on event-related EEG oscillatory activity observed using time-frequency decomposition. Cortical neurons can synchronize their firing with oscillations, allowing neurons participating in the same rhythm to fire with remarkable temporal precision (Gray et al., 1989). This process is crucial for neural firing synchronization and communication through temporal binding, underlying bottom-up and top-down sensory processing (Engel et al., 2001; Lisman & Jensen, 2013; Roux et al., 2022).

In neonates, oscillatory activity and its synchronisation are central to the activity-dependent self-organisation of neural networks, contributing to the stabilization and pruning of synaptic connections essential for cortical development (Ben-Ari, 2001; Hebb, 1949; Khazipov & Luhmann, 2006; Lopes da Silva, 2013; Luhmann & Khazipov, 2018). The emergence of oscillatory activity in the EEG of preterm newborns reflects the progressive maturation of cortical and subcortical neural networks during the late gestational period. In preterm newborns, at term equivalent age, immature EEG discontinuous activity and burst discharges have been progressively replaced by rhythmic oscillatory EEG activities, linked to the maturation of brainstem cortical input (Colonnese et al., 2010; Kostović et al., 1995; Vanhatalo & Kaila, 2006). This process is made possible by the maturation of the properties of interneural network subtending GABAergic transmission (Arshad et al., 2016; Kostović & Judaš, 2002; Vanhatalo & Kaila, 2006) as well as thalamocortical connectivity (Taymourtash et al., 2023), both crucial for the emergence of low and high-frequency oscillatory responses (Buzsáki & Draguhn, 2004; Khazipov et al., 2013). In infants around 28 weeks of gestation, EEG patterns are typically characterized by low-voltage discontinuous activity with prolonged interburst intervals and the presence of delta brushes—slow delta waves (0.5–1.5 Hz) superimposed with faster rhythms (8–25 Hz)—which are considered key markers of early thalamocortical development. Between 28 and 36 weeks gestational age, these patterns evolve toward increased continuity, regional specificity of delta brushes, and the gradual appearance of faster oscillations such as theta and alpha-like rhythms, indicating strengthening of synaptic connectivity and functional organization (Ferrari et al., 1992). The emergence of oscillatory activity in turn has been shown in mice to be crucial for the regulation of interneurons networks and to influence interneural survival as well as the excitation-inhibition balance in the first week post-natal (Duan et al., 2020). Furthermore, sensory stimulus-driven gamma oscillations in the rat mechanistically differed from adult gamma oscillations as they were of thalamic origin and were instrumental in thalamic and cortical barrel maturation as well as thalamocortical functional connectivity (Khazipov et al., 2013; Minlebaev et al., 2011). Consequently, the formation and maturation of cortical and cortical-subcortical networks heavily rely on neuronal activity, with synchronized oscillations involved in stabilizing and pruning neural connections (Hebb, 1949). Thus, synchronized oscillations serve as indicators of cortical network maturity, as they are shaped by anatomical and physiological changes occurring throughout development (Battro & Dehaene, 2010; Buzsáki & Draguhn, 2004).

Event-related oscillatory responses, investigated using time-frequency decomposition, can take the form of an event-related synchronization (i.e. an increase of synchrony in a neuronal ensemble in a given frequency band relative to baseline) or an event-related desynchronization (i.e. a decrease of synchrony in a given frequency band relative to baseline). These oscillations play a causal role in several motor, cognitive and affective processes as shown by experimental non-invasive modulations of these oscillations in adults (Herrmann et al., 2016). Neural synchrony has also been extensively linked to cognitive and perceptual functions, as well as to the development of cortical circuits. It supports synaptic plasticity and reflects ongoing changes in the frequency and coordination of neural activity during maturation (Uhlhaas et al., 2010). Interestingly, in full-term neonates at term-equivalent age, EEG studies employing time-frequency decomposition have shown that infants around 37 weeks of gestational age are able to track speech and singing through low-frequency band activity. These infants also exhibit low-frequency oscillatory patterns capable of discriminating language structures (Ortiz-Barajas et al., 2023) and display a nested oscillatory organization with cross-frequency coupling, reminiscent of patterns observed in adults and non-human primates (Ortiz-Barajas et al., 2023). These findings highlight the relevance of time-frequency decomposition in investigating high-level auditory processing, such as responses to maternal singing, in neonates at term-equivalent age. In this context, examining brain synchronisation across frequency bands offers a valuable tool for assessing brain activity and maturation, particularly in auditory processing in preterm infants.

### Binaural perception of auditory stimuli

Our second goal was to investigate the neural response to binaural stimuli in preterm infants, an area that, to our knowledge, has not yet been explored.

Auditory spatial perception in newborns refers to the ability to detect and localize sound sources in the surrounding environment, a foundational capacity for early social engagement and sensory-motor development. Behavioral evidence suggests that full-term newborns can orient toward lateralized auditory stimuli, indicating the presence of basic spatial hearing mechanisms from birth (Morrongiello et al., 1990). It is a fundamental capacity at the base of early social interactions and sensory stimuli perception (Muir & Field, 1979). This capacity is already present in the first days of life, as full-term newborns exhibit head-turning and eye-orienting responses to lateralized auditory stimuli, suggesting the presence of rudimentary spatial hearing mechanisms (Field et al., 1980). This early sensitivity is believed to rely on basic binaural cues such as interaural time and level differences, processed by subcortical auditory structures that are functional at birth (Moore, 1987). Recent ERP and EEG studies have demonstrated that newborns are neurally sensitive to spatialized auditory stimuli. Notably, brain activity supporting binaural spatialization appears to be present as early as 1–3 days after birth, as evidenced by both early (90 ms) and late (300 ms) ERP responses to spatialized white-noise oddball deviants (Németh et al., 2015). These findings suggest that the neural mechanisms underlying spatial hearing are already functional shortly after birth. Similar ERP responses to spatialized auditory input have also been observed in 4-month-old infants, indicating a developmental continuity in the processing of spatial sound cues (Small et al., 2017). Understanding these interactions is crucial for revealing how early auditory experience shapes the integration of spatial and social auditory cues in infancy. Critically, new findings indicate that this spatial sensitivity extends beyond unimodal processing. In newborns as young as 18 hours old, multisensory integration responses—elicited by concurrent tactile and spatially localized auditory stimulation—showed spatial modulation, with stronger electrophysiological responses to sounds presented near the body compared to those delivered farther away (Ronga et al., 2021). This pattern mirrors adult-like multisensory integration and suggests that neonates possess a primitive, yet functionally relevant representation of the body in space.

However, the time-frequency dynamics of how newborns process spatialized auditory signals, and how these interact with the neural mechanisms underlying voice processing, remain unexplored.

Across a range of EEG studies, a consistent pattern emerges linking binaural spatial sound processing and auditory spatial attention in adults, with distinct oscillatory dynamics, particularly in the beta and alpha frequency bands. Ignatious et al, demonstrated that binaural auditory stimulation modulates high-beta and low-gamma power in midline regions, implicating these frequencies in spatial auditory processing (Ignatious et al., 2023). Complementary findings revealed that frontal-central beta band desynchronization reliably predicts spatial attentional focus, especially when competing sound sources are spatially separated (Geirnaert et al., 2020). Further reinforcing the role of beta oscillations, studies using naturalistic stimuli found increased beta-band involvement during spatialized “Surround” sound as opposed to monaural presentations (Langiulli et al., 2023). At the same time, alpha-band activity has been shown to predict sound source trajectories, as in the tracking of moving auditory stimuli (Bednar & Lalor, 2018) and to lateralize according to attentional demands in dichotic listening paradigms (Kerlin et al., 2010; Wöstmann et al., 2016). In adults, a coherent functional network in which beta oscillations support spatial cue processing, while alpha dynamics mediate attentional orientation within spatial auditory scenes has thus been delineated.

In the present study, we aim to investigate the neural correlates of maternal singing perception in preterm newborns. To address this question, we incorporate a multidimensional approach: (1) examining the effects of familiarity by comparing maternal singing to that of a stranger singing (2) comparing maternal singing to an instrumental rendition of the same melody, thereby introducing a vocal versus non vocal melody contrast; and (3) assessing the influence of auditory delivery methods using binaural context to determine its impact on auditory processing in preterm newborns.

## Materials and Methods

### Population

Twelve preterm infants (mean gestational age at birth: 26.8 weeks), all born at less than 32 weeks of gestational age, were recruited for the current study from the NICU of the Geneva University Hospitals. The clinical characteristics of the infants are presented in Table 1.

**Table 1.**
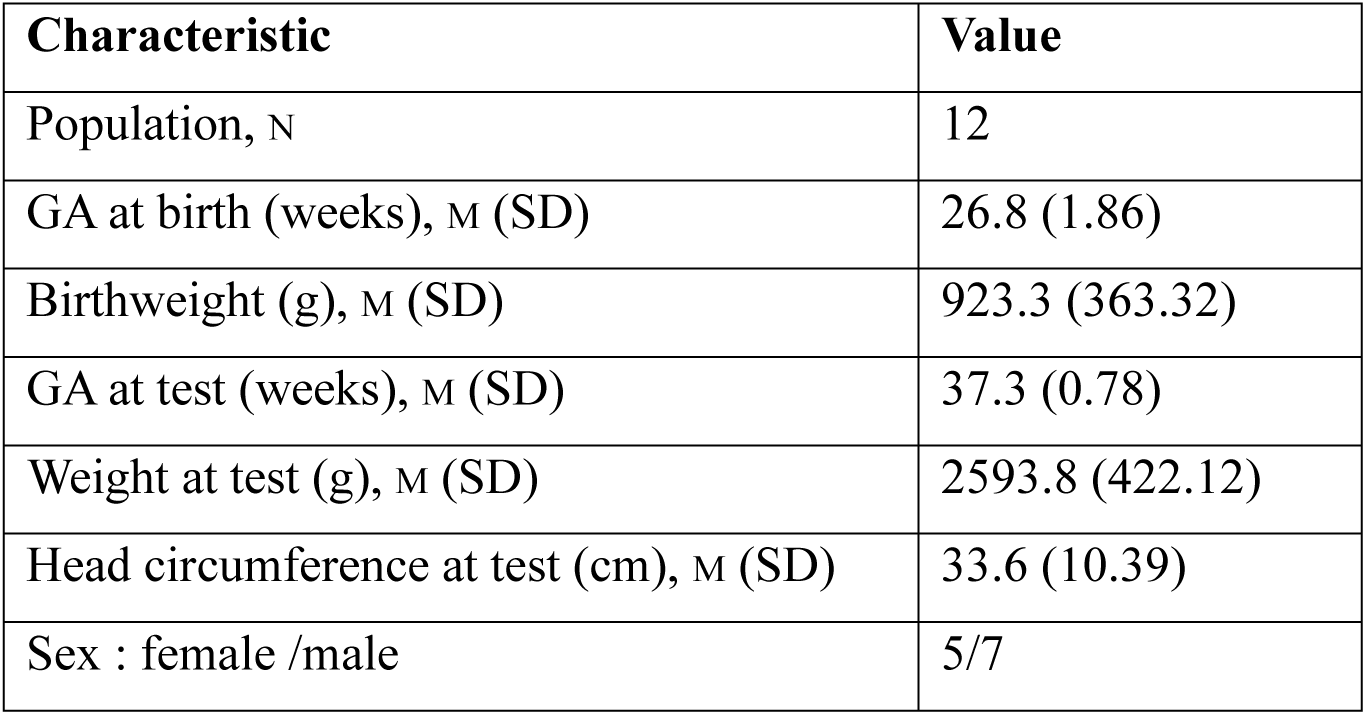
Characteristics of the Population.

Infants with severe neurological complications, such as brain injuries including periventricular leukomalacia and intraventricular hemorrhage, as well as those with hearing or other developmental problems, were excluded from the study. Mothers with history of chronic mental health conditions or alcohol or drug abuse were excluded from the study.

The Swiss Research Ethics Committee approved the study (BASEC no. 2023-00718), and written informed consent was obtained from all parents, in accordance with the Declaration of Helsinki.

### Stimuli and task

Prior to the experiment, the singing voice of each mother was recorded in the NICU using a Neumann M149 microphone and a Grace Design M108 preamplifiers. In the monaural setting, each mother’s voice was placed in the center of the stereo field. For binaural voices, we used a Neumann KU100 artificial head and two Grace Design M108 preamps, and mothers were asked to move in front of the head (always the same movement towards the child, from his right to his left) during the recording, allowing each mother’s voice recording to move in the 3D stereo space in the same way. After EEG installation, the babies were placed in their NICU bed with their parents in the near vicinity. Infants underwent six blocks of auditory stimuli presentation. Each block contained 54 stimuli (duration: 5s), six stimuli per condition, and nine conditions in total. The stimuli were presented in a pseudo-random order, with no possible repetition of the same condition twice in a row. The experimental procedure is represented in Fig. 1.

**Figure 1.**
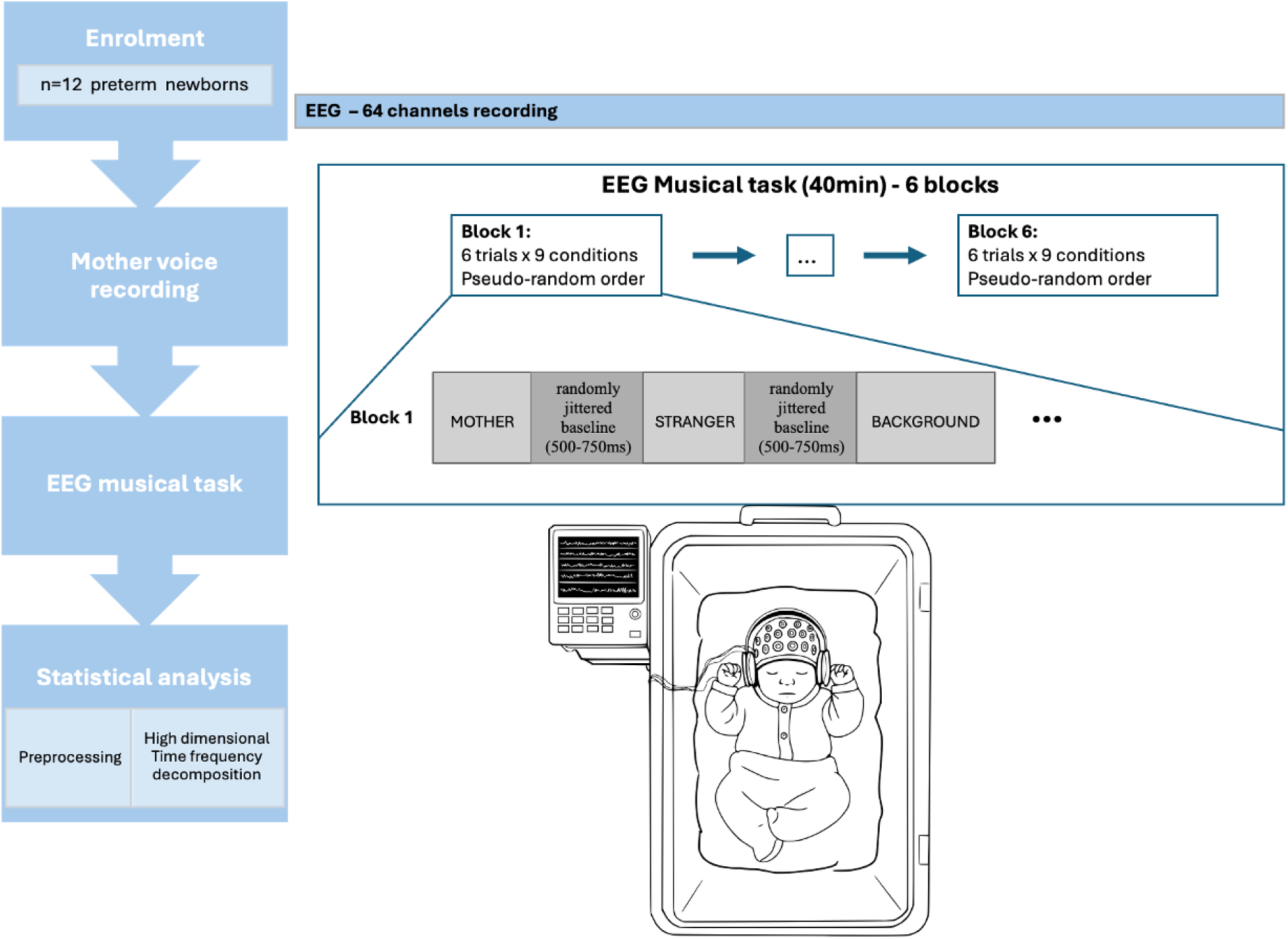
Experimental Procedure. An example of the sequence of auditory stimuli is used to show the experimental procedure. In the study 12 preterm infants were enrolled form Neonatal Intensive Care Unit (NICU) of the Geneva University Hospitals. Each mother voice was recorded in the NICU using a Neumann M149 microphone and a Grace Design M108 preamplifiers. After EEG installation, the babies were placed in their NICU bed with their parents in the near vicinity. EEG was recorded using an ANT Neuro waveguard™ saline-soaked EEG net with a 64-channel. The infants underwent 6 blocks of auditory stimuli presentation. Each block contained 54 stimuli, 6 stimuli per condition, and 9 conditions in total. The stimuli were presented in a pseudo-random order, with no possible repetition of the same condition twice in a row. Statistical EEG analysis was performed using MNE-Python and customs python scripts for preprocessing and following high-dimensional time frequency decomposition.

The auditory stimuli consisted of nine distinct conditions: Background, Mother Voice Monaural and Mother Voice Binaural, Stranger Voice Monaural, Instrument Monaural and Instrument Binaural, Faded Mother Voice Binaural, Gamma Isochronic and Gamma Binaural Beat. Note that each condition contained an auditory background, comprised of harmonic overtones, created from human voices with acoustic synthesis techniques, with a 250ms fade-in at the beginning. The first condition, Background, served as the control. The second condition, Mother Voice Monaural, featured the infant’s mother singing two notes superimposed on the background, where the mother’s voice began 900 ms after the onset of the background and the note change occurred at 1.65 s. The notes sung or played were the major third of D (F sharp) and the fifth of D (A). Similarly, the third condition, Stranger Voice Monaural, consisted of the singing voice of another mother enrolled in the study, following the same temporal structure as the Mother Voice Monaural. In the fourth condition, Mother Voice Binaural, the mother’s voice was spatialized binaurally while maintaining the same temporal structure as the monaural conditions. The fifth condition, Instrument Monaural involved the background combined with a punji instrument playing the same notes as in the vocal conditions, and the sixth condition, Instrument Binaural, utilized the same notes as the Instrument Monaural condition but spatialized binaurally. These vocal stimuli were matched with vocal stimuli for intensity and pitch. Finally, the seventh condition, Faded Mother Voice Binaural, incorporated the infant’s mother’s singing voice faded into the background. In the present study, we focused on describing and analyzing seven of the nine experimental conditions, excluding Gamma Isochronic and Gamma Binaural Beat Stimuli from the current analysis. A pictural description of these seven stimuli are listed in Fig 2.

**Figure 2.**
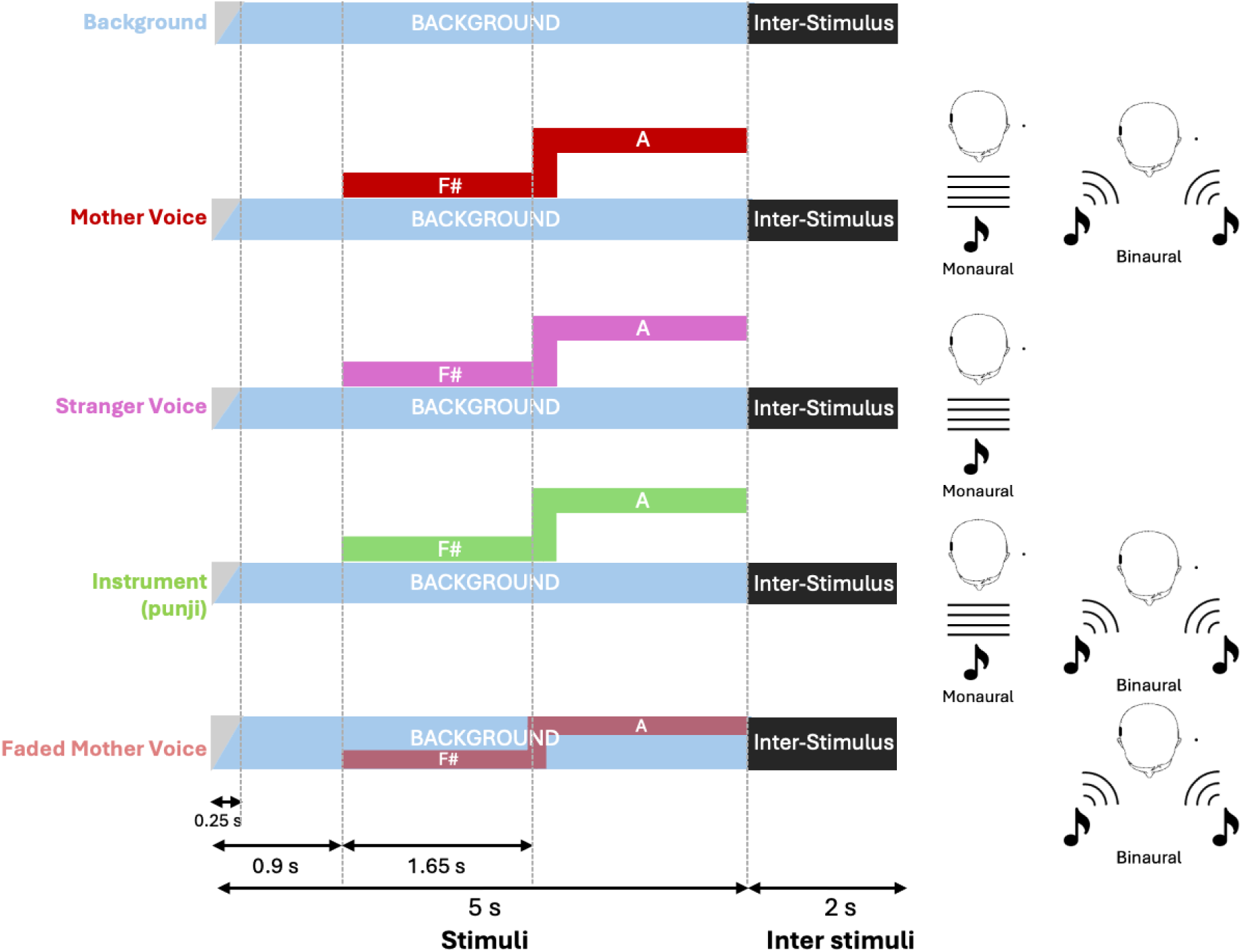
Pictural description of the seven auditory conditions. Seven of the nine auditory conditions which were presented to preterm infants, are discussed and analyzed here. In each condition, the auditory stimulus began 900 ms after the onset of the background, and a note change occurred at 1.65 seconds. The musical notes used were the major third (F♯) and the fifth (A) of the D major scale. Vocal and instrumental stimuli were carefully matched for both intensity and pitch.

Trial structure was as follow: following a randomly jittered silence (500-750ms), the stimuli was presented for 5s, followed by a 2s inter-stimuli interval.

The first condition, *Background*, consisted of harmonic overtones synthesized from human voices and presented as a stereophonic sound. In the *Mother Voice Monaural* condition, the infant’s mother’s singing voice was added to the background. The *Stranger Voice Monaural* condition included the background with the singing voice of another mother enrolled in the study. In the *Mother Voice Binaural* condition, the mother’s voice was spatialized binaurally while maintaining the same temporal structure as in the monaural condition. The *Instrument Monaural* condition combined the background with a punji instrument playing the same notes as in the vocal conditions. The *Instrument Binaural* condition used the same instrumental notes as the monaural version but was spatialized binaurally. Finally, in the *Faded Mother Voice Binaural* condition, the mother’s singing voice was progressively faded into the background and presented binaurally.

The trial structure was as follows: after a randomly jittered silent baseline (500–750 ms), the stimulus was presented for 5 seconds, followed by a 2-second interstimulus interval. The background, composed of harmonic overtones derived from human voices and beginning with a 0.25-second fade-in, is present across all conditions. A silent inter-stimuli interval of 2 seconds follow each stimulus. In red, the mother’s voice sings the note F# starting 0.9 seconds after stimulus onset for 1.65 seconds, followed by the note A until the end of the stimulus. In pink, the stranger’s voice performs the same notes. In green, the punji (a traditional Indian flute) plays the identical notes. In faded red, the mother’s voice is incorporated as a background component. The different musical stimuli were carefully matched for intensity and pitch to ensure comparability.

### EEG recordings

The EEG was recorded throughout the task using an ANT Neuro waveguard™ saline-soaked EEG net with a 64-channel equidistant hexagonal layout, connected to an eego™ amplifier. The setup was optimized to ensure high-quality signal acquisition, enabling comprehensive analysis of neural responses during the auditory stimuli presentation.

The net was secured using a tissue cap. The data were acquired at a sampling rate of 1000 Hz. Electrode impedances were typically maintained below 50 kΩ. The infants’ state was coded prior to and during each experimental block by a trained researcher. Data were collected while the newborns were either awake or in a state of sleep, quiet or deep, without sedation. To reduce agitation during the task, the newborns were fed prior to testing. The infant’s parents were present in the room and could provide soothing pressure on their child if agitated.

### EEG data processing and statistical analysis

EEG analysis was performed using MNE-Python and customs python scripts (Gramfort, 2013; Larson et al., 2024). Preprocessing steps were applied to raw EEG recordings (down-sampling at 250Hz, bad channels interpolation, average referencing, fir filtering with hamming windows from 0.5 to 120Hz, notch filtering of line-noise at 50Hz). Epochs were cut from -4 to 7s post-sound onset, to avoid border effects for time-frequency computing, heavily artifacted periods were rejected (peak to peak amplitude for any channel above 600uV), and artifacts due to eye movements or heart beats were rejected using ICA.

Multitaper time-frequency decomposition was performed from 2 to 100Hz in 1Hz steps using 1 slepian taper and a 500ms window for low-frequencies(<40Hz), and 5 slepian tapers and a 200ms windows for high frequencies (>40Hz) from -0.5 to 5s in 80ms steps (70 bins, Ferrante et al., 2022). Time-frequency data was converted to dB and baseline corrected for -500 to - 100ms pre-sound onset, to ensure onset activity contamination avoidance, as preliminary analysis suggested spectral gamma-leakage of onset-activity in the -100 to 0ms time-interval, due to temporal smoothing.

To perform spectral outlier rejection, physiological range in low and high broadband spectrum was assessed by calculating, for all participants, channels and trials altogether the average power in the 2-12Hz, 12-30Hz, 30-45Hz and 55-100Hz frequency range. For each participant, in each channel, trials where average power in any of these 4-frequency range fell outside the 95% confidence interval (average +- 1.96std of the whole trial/channels/participants distribution) for the frequency range considered was rejected for subsequent analysis.

Data were analyzed using large scalp ROIs (see SI_Fig4, Frontal, Central, Right Temporal, Left Temporal, Occipital) as well as a more precise topographical analysis of effects (see below).

A first analysis aimed to investigate the precise time course of the time-frequency effects of soundscape elements presentation in large-scalp ROI. For each condition, in each ROI, time-frequency maps for each trial in each channel in the ROI were averaged. For each trial, in each channel, at each time points, the average power was calculated in 11 frequency bands (delta: 2-4Hz, theta: 4-8Hz, alpha: 8-12Hz, beta12: 12-20Hz, beta20: 20-30Hz, gamma30: 30-45Hz, gamma55: 55-60Hz, gamma60: 60-70Hz, gamma70: 70-80Hz, gamma80: 80-90Hz, gamma90: 90-100Hz) and entered a database.

Then, for each ROI, for each frequency band, at each timepoint, data is entered into General Linear Mixed models testing the main effect of Condition (10 levels) on Power with the ID and Channel as random factors. If a main effect of Condition is found, contrast analyses are performed to test several hypotheses: 1. the effect on oscillatory activity of the Mother Voice and Stranger Voice in a monaural setting (Condition: 3 levels: Background, Mother Voice Monaural and Stranger Voice Monaural); 2. the effect on oscillatory activity of Binaural versus Monaural presentation of Vocal and Instrumental stimuli (Condition: 4 levels: Mother Voice in the Monaural and Binaural setting, Instrument in the Monaural and Binaural setting); 3. the effect on oscillatory activity of Faded Mother compared to the background (Condition: 4 levels: Background, Mother Voice in the Monaural and Binaural setting, Faded mother voice). For each hypothesis, if the effect of Condition reached significance (p<0.05, corrected for multiple comparisons), contrasts between conditions and effects size were calculated using the emmeans package. P-values were false-discovery rate corrected (Benjamini & Hochberg, 1995).

A second analysis aimed at investigating more precisely the topographical time course of the time-frequency effects of soundscape elements presentation. To this aim, 19 smaller scalp ROIs were constituted (14 channels triplets, 4 channels doublets, and one occipital cluster with 5 channels, SI Fig 1). A similar statistical analysis was performed as described in the previous paragraph and p-values were corrected for all comparisons in the whole cluster pool using the FDR method.

## Results

### Maternal singing in the monaural setting elicits a low-frequency whole-scalp synchronisation

Preterm newborns exhibited distinct patterns of brain synchronisation across frequency bands, scalp topographies, and temporal windows in response to the same melody when sung by the mother, a stranger, or played by an instrument.

Figure 3 presents the baseline corrected time-frequency maps for the background, mother voice, and stranger voice in the monaural setting (Figure 3.1), as well as contrasts between conditions (Figures 3.2 and 3.3) and topography of the mother versus stranger voice effect (Figure 3.4). The frontal cluster has been selected for representation as it presented the earliest significant contrasts and with the highest amplitude.

**Figure 3:**
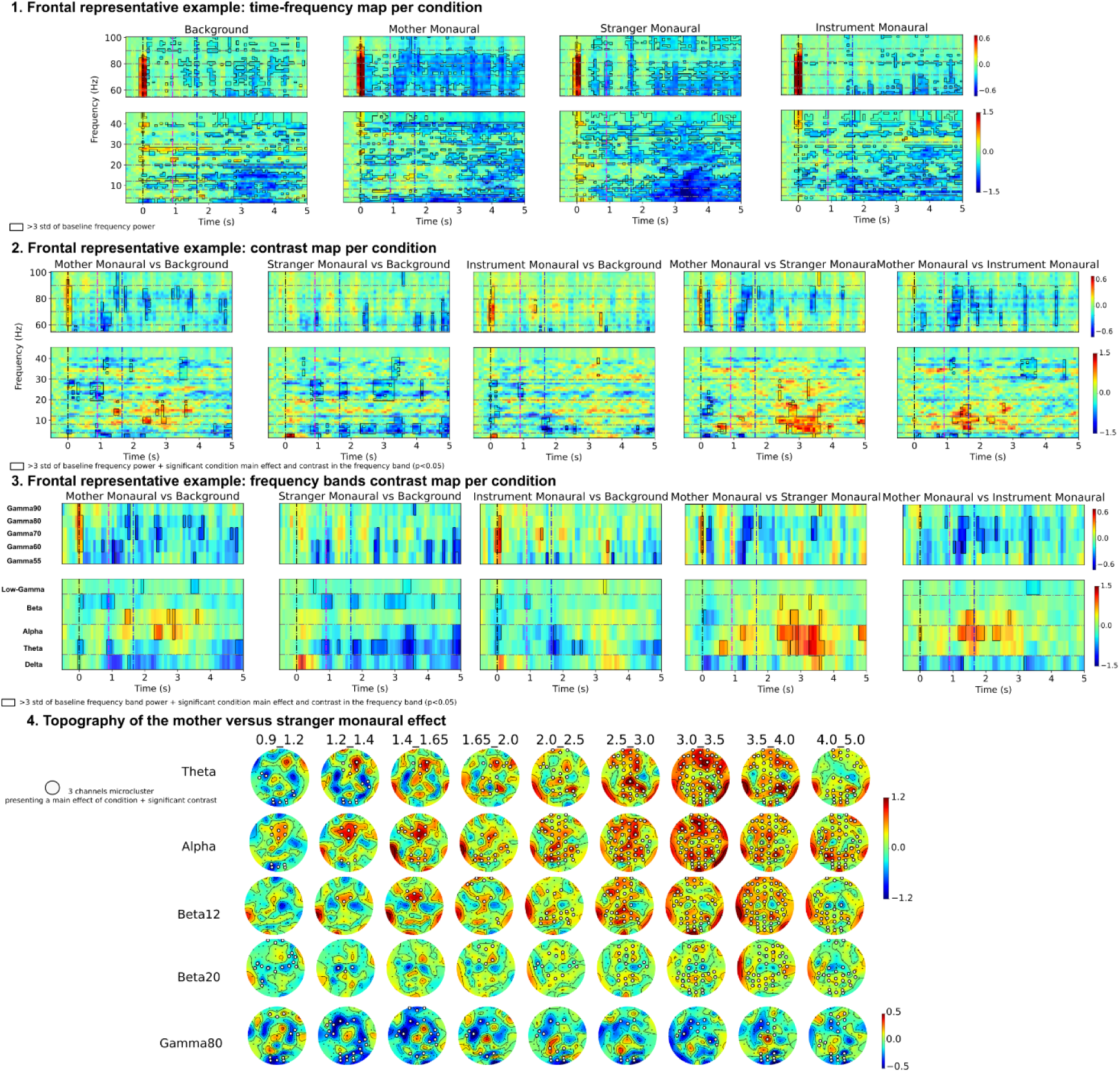
Preterm infants’ EEG responses to the monaural presentation of mother voice, stranger voice and instrument. 1. Baseline corrected time-frequency maps for background, mother voice, and stranger voice in the monaural setting, framed areas represent time and frequencies where absolute baseline corrected power exceeds 3 times the standard deviation of mean baseline power. 2. Contrasts between conditions, all frequencies are represented at each timepoint. 3.Simplified representation of contrasts between condition with the average power for each frequency band statistically investigated. Framed areas represent time and frequencies where **3** statistical criteria are satisfied: First, absolute baseline corrected power exceeds 3 times the standard deviation of mean baseline power; Second, GLM analysis yielded a significant effect of condition for this frequency band at this timepoint and third, contrast analysis yielded a significant effect of the pairwise contrast considered after FDR correction. For 1-2-3, the frontal cluster has been selected for representation as it presented significant contrasts with the highest amplitude. 4.Scalp topography of the mother versus stranger effect. White dots indicate channels within a 3-channel micro-cluster where the GLM analysis revealed a significant effect of condition for the given frequency band and time point. Additionally, the pairwise contrast analysis showed a significant effect after FDR correction, offering a more precise topographical localization of the observed effects. FDR: False Discovery Rate, GLM: General Linear Mixed.

The main Results are summarized in Table 2.

**Table 2.**
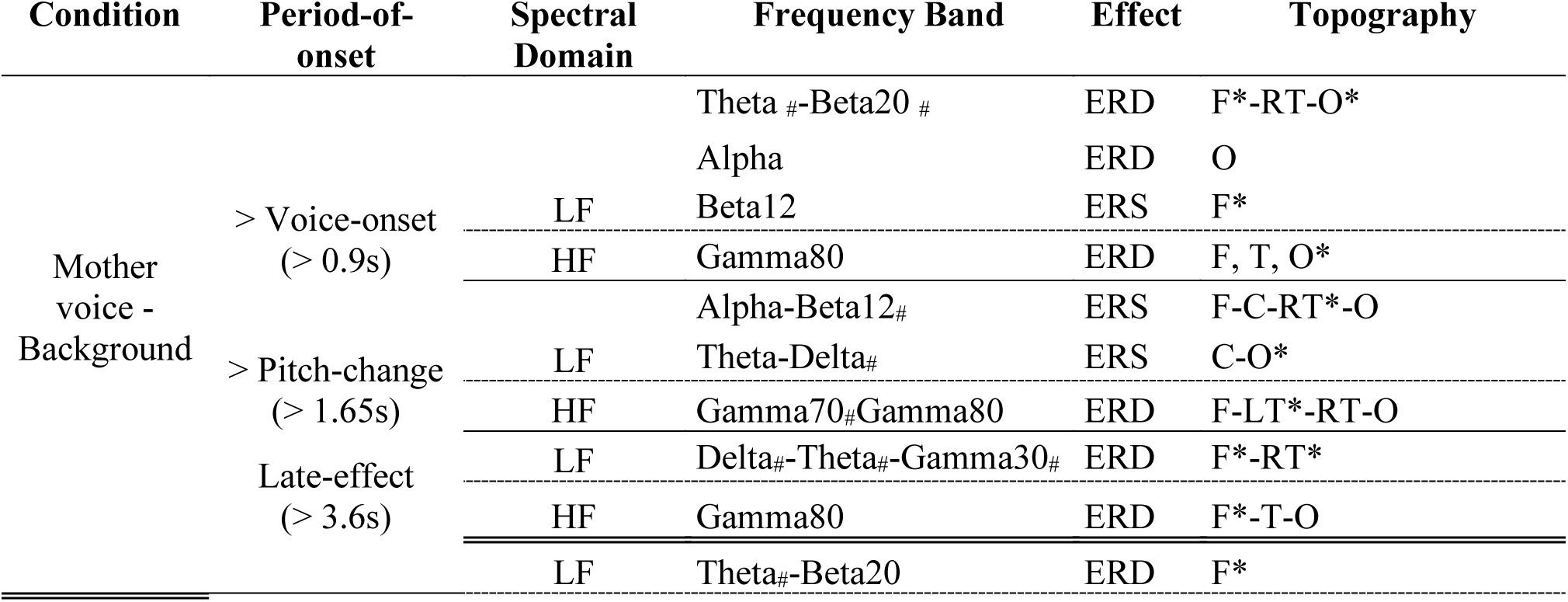
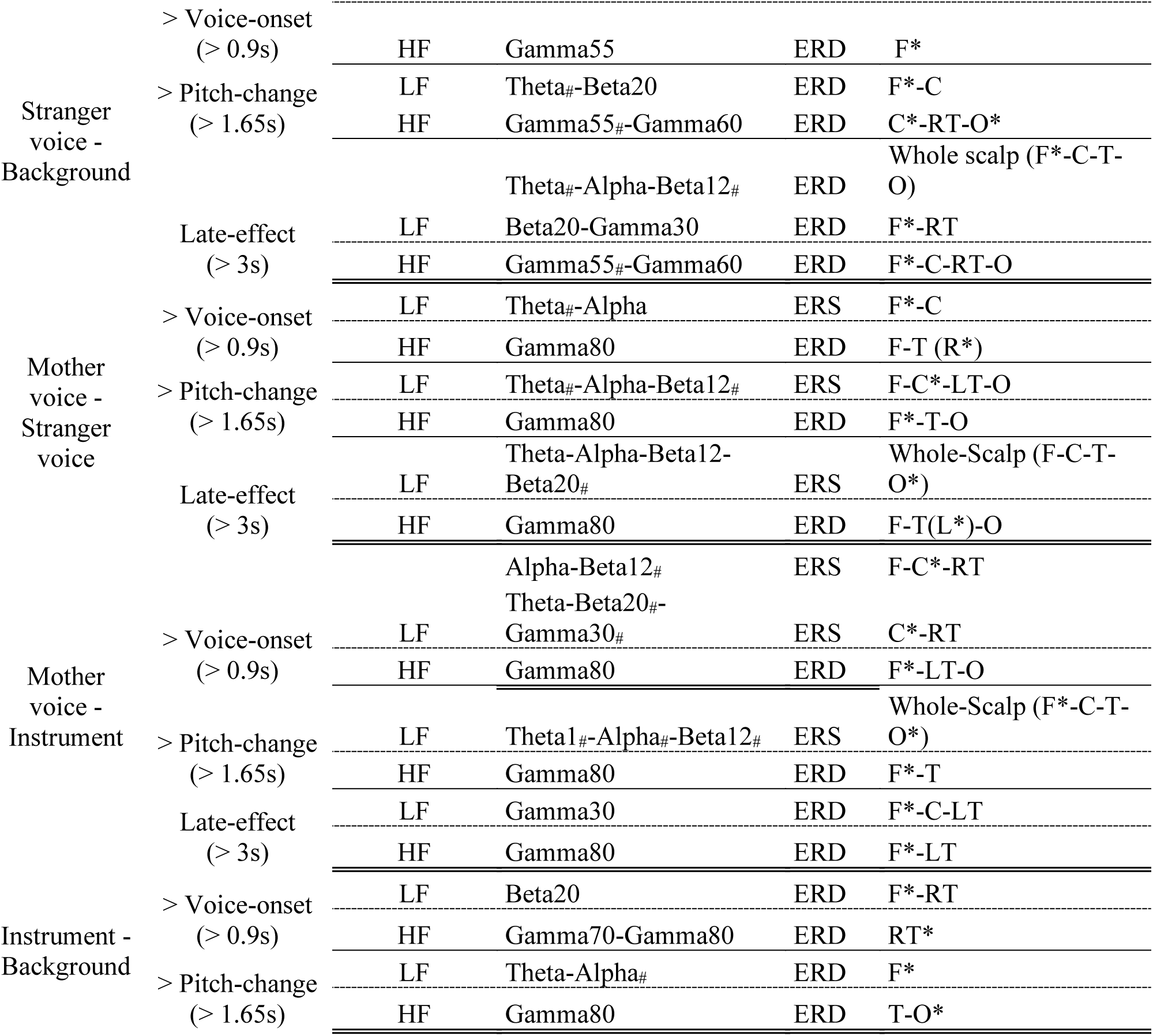
Summary of the main EEG Contrasts in response to monaural presentation of mother voice, stranger voice, and instrument in premature infants. In the first columns the contrasts of interest are presented, and in the second column the main time period when the reported effects occur. Spectral-domain: low-frequencies (LF, delta, theta, alpha, beta12, beta20, and gamma30) and high-frequencies (HF, broadband gamma: Gamma55, Gamma60, Gamma70, Gamma80, Gamma90). Frequency band; Delta:2-4Hz, Theta: 4-8Hz, Alpha: 8-12Hz, Beta12: 12-20Hz, Beta20: 20-30Hz, Gamma30: 30-45Hz, Gamma55: 55-60Hz, Gamma60: 60-70Hz, Gamma70: 70-80Hz, Gamma80: 80-90Hz, Gamma90: 90-100Hz. Effect: event-related synchronization (ERS) or event-related desynchronization (ERD). Topography: spatial ROI clusters where the reported effects reached significance (C=Central, F=Frontal, O=Occipital, T=Temporal right and left, RT=Right temporal, LT= Left temporal). * reports the topographical cluster where the reported effect occurs the earliest. <u>Note</u>: in the frequency band columns, # materialize frequency bands where the effect reported did not reach significance for all clusters in the topography. A graphical representation of contrasts for all topographical clusters is available in SI Fig 1. Only major contrasts are presented in this table, minor significant effects are also available in SI.

### Background and Mother Singing Condition

Overall, a whole-scalp low-frequency (theta, alpha, and beta bands) event-related desynchronization (ERD) relative to baseline (3* baseline power standard deviation) was mainly observed in response to background overtones and monaural mother voice presentation except from voice-onset to 3s in the theta-alpha-beta12 bands for the monaural mother-voice condition (Figure 3). After a transient theta-beta20 ERD and beta12 event-related synchronization (ERS) around sound onset most consistent in the frontal regions, a significant relative ERS was observed for the monaural mother voice compared to the background in the post-pitch change (>1.65s) period in the theta, alpha, beta12, beta20, and occipital delta (Table 2).

In the broad gamma bands, a whole scalp gamma ERD was observed for background and mother monaural presentation (Table 2). A significantly stronger frontal-temporal-occipital ERD was observed for the monaural mother voice compared to the background, most consistently in the gamma80 band (Table 2).

### Stranger Singing Condition

Following stranger voice presentation, a whole-scalp low-frequency ERD is also observed relative to baseline (starting around sound-onset). A frontal-central relative ERD for stranger voice compared to background was observed in the theta and beta20 bands starting around voice onset. This ERD effect extended to broadband gamma55 after pitch change in frontal-central-occipital areas and beta12/alpha bands in the whole scalp 3s post-pitch change. Late Post Stimulus Onset (PSO) (>3s PSO) relative ERD compared to the background was also observed in the delta and beta20/gamma30 bands (Table 2).

Consequently, mother voice compared to the stranger voice presented a whole scalp low-frequency ERS (theta-alpha-beta12 bands, Table 2), occurring earliest after voice presentation in frontal-central areas (>0.9s PSO), as well as a consistent frontal-temporal gamma70-80 relative ERD occurring 1.2s PSO.

### Instrumental Condition

The effect described above was specific to vocal stimuli. Indeed, the monaural punji-instrument control mainly elicited a frontal-temporal early (>0.9s) beta20 ERD followed by later (>1.65s PSO) theta-alpha band ERD relative to the background. Mother voice compared to instrument elicited a low-frequency whole scalp relative theta-alpha-beta12-beta20-gamma30 ERS (Table 2) as well as a broadband gamma80 relative ERD starting around 1.2s PSO after voice onset.

To summarize, maternal voice in the monaural condition elicited a low-frequency ERS and gamma80 long-lasting ERD compared to baseline, background, stranger monaural, and instrument monaural conditions. The most consistent contrast across conditions occurred in frontal-central areas (>1.65s PSO).

### Binaural presentation of musical stimuli elicits a low-frequency event-related desynchronisation across voice and instrumental modality

We then aimed to examine the effect of binaural spatialization on the oscillatory processing of the mother voice and instrument in our preterm population. Baseline corrected time-frequency maps for the monaural and binaurally presented mother voice and instrument are represented in Figure 4.1, and contrasts between conditions are represented in Figure 4.2 and 4.3, and topography of the mother voice monaural versus binaural effect is represented in Figure 4.4. The frontal cluster has been selected for representation for comparison purposes with monaural contrast. The main Results are summarized in Table 3. Other spatial clusters are presented in SI Fig 4.5.

**Figure 4:**
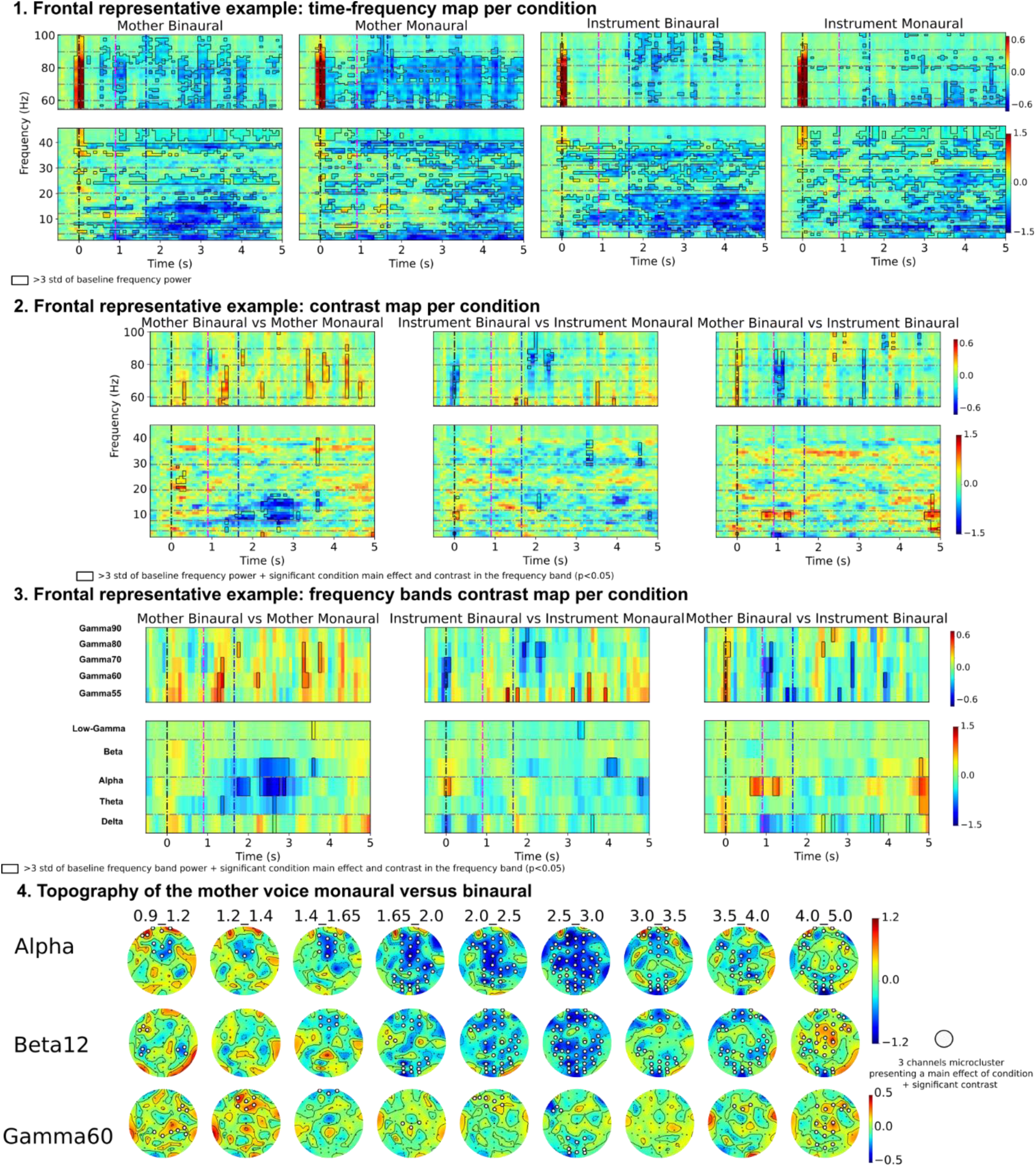
Preterm infant’s EEG response to the binaural presentation of mother voice and instrument. 1. Baseline corrected time-frequency maps for monaural and binaural presentation of mother voice and instrument, framed areas represent time and frequencies where absolute baseline corrected power exceeds 3 times the standard deviation of mean baseline power. 2.Contrasts between conditions, all frequencies are represented at each timepoint. 3. Simplified representation of contrasts between conditions with the average power for each frequency band statistically investigated. Framed areas represent time and frequencies where 3 statistical criteria are satisfied: First, absolute baseline corrected power exceeds 3 times the standard deviation of mean baseline power; Second, GLM analysis yielded a significant effect of Condition for this frequency band at this timepoint and third, contrast analysis yielded a significant effect of the pairwise contrast considered after FDR correction. For 1. 2. 3. the frontal cluster has been selected for representation to allow for easier comparison with the monaural mother versus stranger effects. 4. Scalp topography of the mother voice monaural versus binaural effect. White dots indicate channels within a 3-channel micro-cluster where the GLM analysis revealed a significant effect of condition for the given frequency band and time point. Additionally, the pairwise contrast analysis showed a significant effect after FDR correction, offering a more precise topographical localization of the observed effects. FDR: False Discovery Rate, GLM: General Linear Mixed.

**Table 3:**
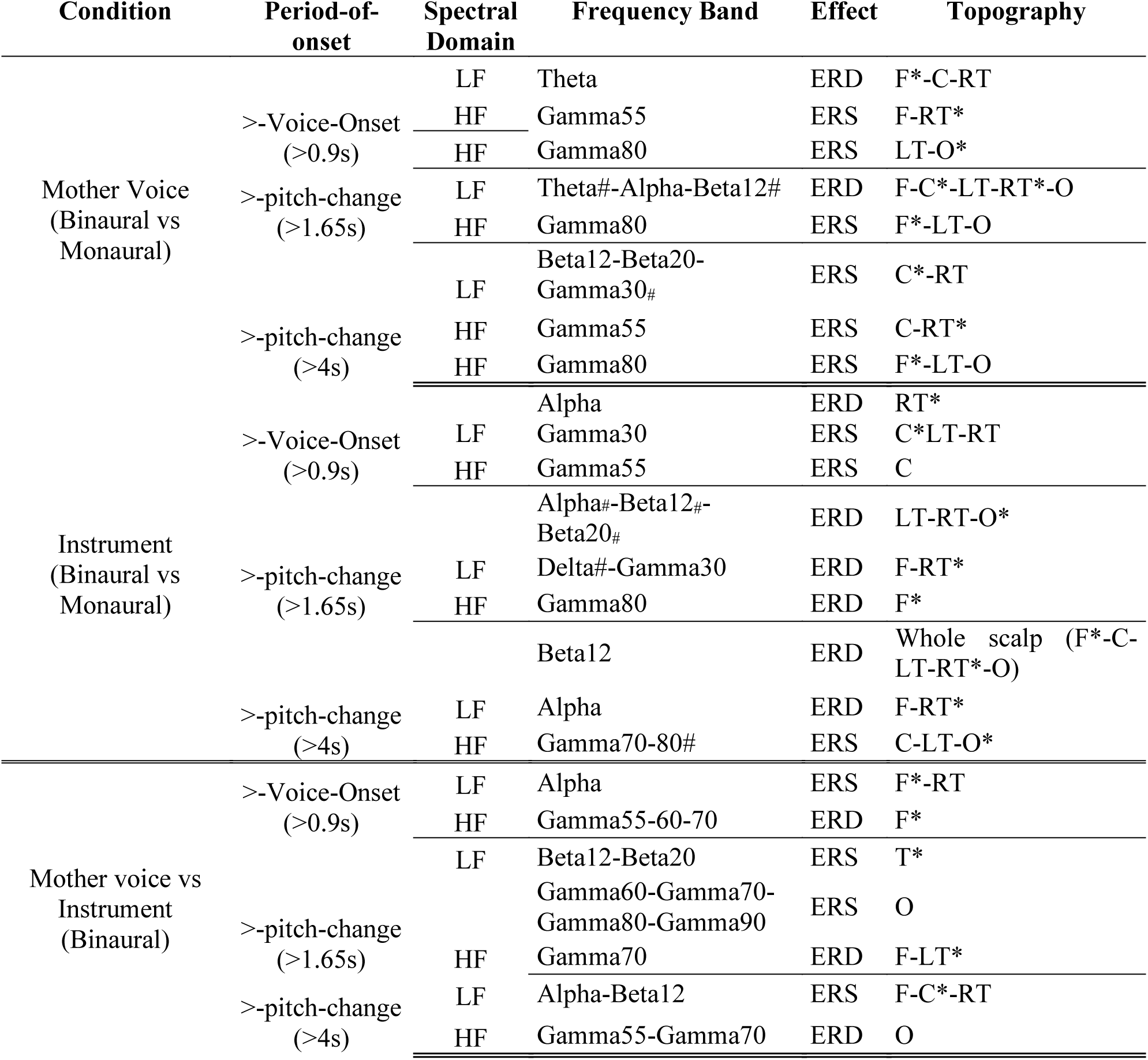
Summary of the main EEG Contrasts in response to monaural versus binaural presentation for mother voice, stranger voice and instrument in premature infants. In the first column are presented the contrasts of interest, and in the second column the main time period when the reported effects occur. Spectral-domain: low-frequencies (LF, delta, theta, alpha, beta12, beta20, and gamma30) and high-frequencies (HF, broadband gamma: Gamma55, Gamma60, Gamma70, Gamma80, Gamma90). Frequency band; Delta:2-4Hz, Theta: 4-8Hz, Alpha: 8-12Hz, Beta12: 12-20Hz, Beta20: 20-30Hz, Gamma30: 30-45Hz, Gamma55: 55-60Hz, Gamma60:60-70Hz, Gamma70: 70-80Hz, Gamma80: 80-90Hz, Gamma90: 90-100Hz. Effect: event-related synchronization (ERS) or event-related desynchronization (ERD). Topography: spatial ROI clusters where the reported effects reached significance (C=Central, F=Frontal, O=Occipital, T=Temporal right and left, RT=Right temporal, LT= Left temporal). * reports the topographical cluster where the reported effect occurs the earliest. <u>Note</u>: in the frequency band columns, # materialize frequency bands where the effect reported did not reach significance for all clusters in the topography. A graphical representation of contrasts for all topographical clusters is available in SI Fig 3. Only major contrasts are presented in this table, all significant effects are available in SI.

### Mother Singing Condition

In response to the binaural mother voice, a strong low-frequency ERD was observed compared to baseline. Hence a whole-scalp low-frequency relative ERD was observed between binaural and monaural mother voice, first (>1.2s PSO) in the theta band, then (>1.65s PSO) in the alpha, and beta12 bands. An early (>1s PSO) mainly frontal-temporal-occipital broadband gamma relative ERS was also observed between binaural and monaural-mother-voice, especially in the gamma55, gamma60 and gamma80 bands (Table 3).

### Instrumental Music Condition

Similarly, binaural instrumental stimuli elicited a low-frequency whole scalp ERD relative to baseline. Hence, a low-frequency relative ERD was observed post-pitch-change (>1.65s) between binaural and monaural-instrument in the alpha, beta12, beta20, and gamma30 bands across scalp ROIs (Table 3). In the broadband gamma bands, a biphasic pattern was observed: first, a relative frontal-temporal gamma80 ERD for binaural instrument presentation compared to monaural followed by a mainly central-occipital relative gamma80 ERS after 4s PSO.

Note that despite the concurrent desynchronization in both the mother-voice and instrument modality, the binaural mother-voice compared to instrument still yielded a significant mainly temporal and frontal post-voice onset alpha-beta12 relative ERS, as well as a relative occipital broadband gamma ERS post-pitch change.

### Faded Mother Singing Condition

To test whether preterm neonates could process their mother’s voice and binaurality regardless of dB presentation, the faded mother stimuli were designed where the binaural mother’s voice was faded into the vocal harmonic background. The faded binaural mother pattern of oscillatory activation was highly reminiscent of what was observed for the binaural mother voice. Indeed, as observed for binaural mother voice stimuli, faded mother voice elicited relative to background a relative ERD in the theta (>0.9s PSO), alpha (>1.9s PSO), and beta20 bands (>0.9s PSO, SI-Fig5-6). These results are discussed in more detail in SI.

To summarize, the binaural presentation of maternal voice and instrument elicited a whole-scalp low-frequency ERD relative to their monaural analogs. The mother voice binaural also elicited a relative broadband gamma ERS compared to the monaural setting, while the binaural presentation of instrumental sounds elicited a broadband gamma ERD followed by a late rebound. The relative mother voice versus instrument ERS was reproduced in the binaural setting.

## Discussion

In the present study, we demonstrated that preterm newborns display different patterns of neural responses when the same melody is delivered by the mother, a stranger, or an instrument. Specifically, we identified a unique whole-scalp brain synchronization in preterm newborns during maternal singing processing, over stranger singing or instrumental melody. Infant-directed singing is a unique human behavior (Corbeil et al., 2013; Nakata & Trehub, 2004; Trainor & Cirelli, 2015; Tsang et al., 2017), universally recognizable by its recurring acoustic patterns (Hilton et al., 2022) and serving specific functions during the early stages of infant development (Cirelli et al., 2020; Corbeil et al., 2013, 2016; Lense et al., 2022; Leong et al., 2017; Nguyen et al., 2023; Punamäki et al., 2024; Shenfield et al., 2003).

The present results align with the distinct preterm infants’ behavioral responses to maternal singing, as compared to speech, reported in the literature (Filippa & Kuhn, 2024). Unlike speech, singing helps preterm newborns maintain their current sleep state rather than inducing wakefulness and elicits rhythmical mouth movements that align with the singing’s rhythmicity (Filippa et al., 2020). Notably, it induces a more mature autonomic nervous system response, highlighting its role in supporting physiological regulation (Filippa et al., 2024).

In the present study, we observed a relative whole-brain synchronization in response to the presentation of the mother’s singing voice, particularly in the theta, alpha, and beta bands, with activity concentrated in the frontal, occipital, and temporal regions. In contrast, the stranger’s singing voice elicited desynchronization across multiple frequency bands (theta, alpha, beta, and gamma) in the same scalp regions, especially during early post-stimulus onset.

Theta and delta band activity has indeed been shown to track vocal stimuli (Attaheri, Choisdealbha, et al., 2022; Attaheri, Panayiotou, et al., 2022), even at the newborn stage (Ortiz-Barajas et al., 2023). In adults, such brain-stimulus tracking has been shown in MEG and EEG studies to be to be enhanced by attentional processes (Destoky et al., 2019; Kerlin et al., 2010; Schüller et al., 2025). In the present cohort of preterm infants, attentional processes may also enhance song tracking in the low-frequency bands, potentially accounting for the observed increase in power. Conversely, the diminished synchronization during the presentation of the stranger’s singing voice may reflect reduced attentional engagement, likely due to its lower emotional salience compared to the maternal voice. Interestingly, previous studies investigating older infants’ responses to nursery rhymes (Attaheri, Choisdealbha, et al., 2022; Attaheri, Panayiotou, et al., 2022) did not report significant tracking in the alpha band for infants, possibly because the auditory stimuli used included speech and words, even if embedded in a rhythmical and musical structure. This contrasts with the current findings showing alpha-beta synchronization, suggesting that non-verbal and song-specific features may engage different neural mechanisms in early auditory processing. However, in a study on full-term newborns, a frontal power increase in the theta, alpha and low-beta band was observed during presentation of speech compared to a baseline, in which seems to indicate an effect of voice on alpha power irrespective of tracking processes (Filippa et al., 2023). A low-frequency event-related synchronization during music listening was also observed in children aged 1-12 years (Bower et al., 2024). The synchronization observed in these frequency bands might then represent an attentional modulation of these higher-order voice-related processes beyond low-level voice-envelope tracking. Of note, a phase-amplitude coupling between theta and beta-band activity has been observed during nursery rhymes processing by 4-month-old infants, which might also play a role in the beta synchronization response reported in the current study (Attaheri, Choisdealbha, et al., 2022).

Along with attentional and vocal-perceptive processes, we hypothesize that affective and emotional factors might also play a role in the results reported in the current study. Indeed, the adult literature have reported a correlation between alpha band synchronization and comfort in resting-states EEG studies in adults (Tarailis et al., 2022), as well as an alpha-band power increase after musical training designed to induce relaxation (Phneah & Nisar, 2017). A frontal-posterior-temporal theta-alpha power increase was also observed in adults in the “blissful state” observed during meditation (Aftanas & Golocheikine, 2001; Lagopoulos et al., 2009). Within this framework, we could hypothesize that the synchronization observed during the presentation of the mother’s singing voice may be associated with a ‘soothing effect’ of maternal singing on preterm newborns. This hypothesis is further supported by the timing of the effects observed in the current study. The low-frequency ERS linked to mother-singing presentation occurs relatively late during the trial, consistent with higher order top-down processes, which is to be expected if this activity reflects an affective process linked to mother’s voice recognition. As theta-band synchronization has been observed during affective regulation, our results are altogether in line with a potential beneficial effect of maternal-singing presentation in the adverse setting of the NICU (Takács et al., 2024).

Alternatively, as preventing beta desynchronization has been associated with impaired memory encoding (Hanslmayr et al., 2016), our results might also suggest that directed singing from strangers are processed as novel information requiring new memory encoding (Griffiths et al., 2021).

Importantly, the present results on maternal singing contrast with a previous research on the preterm newborns’ maternal speech perception conduced with similar analyses, where preterm infants showed selective EEG low-frequency responses to stranger voices in both temporal hemispheres, while lacking distinct brain responses to their mother’s voice in the same frequency bands (Adam-Darque et al., 2020; Filippa et al., 2023). The low-frequency response observed in the current study in response to maternal singing might then suggest a differential processing of maternal voice in preterm neonates when presented in sung form compared to speech, highlighting the former as a potentially preserved channel of mother-preterm newborn interaction. In the NICU, preterm newborns are seldom exposed to singing by strangers, whereas they are regularly exposed to stranger speech, particularly from nurses and medical staff.

Moreover, we observed desynchronization in the broadband gamma bands occurring earlier for the mother-voice compared to the stranger-voice. The effect of gamma desynchronization is not yet fully understood, but adults studies suggest its potential role in complex processes in language related areas (Ihara et al., 2003). A computational model of the auditory pathway observed a decrease of power in the gamma band of the neurons responsible for processing the sound input (Uriarte et al., 2016). A systematic review suggests that individual variations in gamma-range auditory steady-state response reflect the capacity for attentional control and the ability to temporarily store and manipulate information—skills essential for complex cognitive activities, including language (Parciauskaite et al., 2021). Alternatively, this gamma ERD response to voices might be atypical manifestation specific to preterm infants during voice-processing. Indeed, in a previous study on maternal speech perception by preterm and full-term newborns (Adam-Darque et al., 2020; Filippa et al., 2023), a gamma ERS was observed in full-terms during mother’s voice presentation while gamma activity in preterm failed to reach significance during presentation of a mother or a stranger voice alike. Furthermore, a qualitative examination of the gamma time-frequency pattern in this previous study suggested, in preterm infants, a gamma desynchronization pattern during mother and stranger-voice presentation presenting striking similarities with the pattern observed in the current study, while no such gamma ERD is observed in full-terms. As early gamma activity, has been shown to be crucial to the organization of thalamo-cortical connectivity, which has been shown to be impaired in preterm infants (Khazipov et al., 2013; Minlebaev et al., 2011), further studies directly comparing gamma activities during maternal singing in preterm and full-term would be of crucial interest to investigate the mechanistic impact of this processes.

Furthermore, in the present study, the instrumental melody evoked limited relative ERS, primarily in occipital areas for alpha and beta bands. This suggests less direct engagement of auditory-specific neural networks compared to vocal stimuli. These results correlate with Loukas et al. results on functional magnetic resonance imaging performed on newborns (full-term and preterm) while listening to melodies played by an instrument or sung by a female voice. Interestingly, vocal stimulus elicited sensorimotor response and was processed as a more salient stimuli while the instrumental condition activated higher-order cognitive in particular the default mode network and visuo-spatial networks (Loukas et al., 2024). The significant alpha and beta ERS for maternal voices versus instrumental sounds, along with gamma ERD, underscored the specificity of maternal vocal stimuli in engaging neonatal auditory and emotional circuits. These results support the hypothesis that maternal singing holds a unique capacity to engage neonatal auditory and emotional circuits, which may have implications for interventions in populations with atypical developmental trajectories, such as preterm infants (Filippa et al., 2017; Haslbeck et al., 2023; Kostilainen et al., 2021; Partanen et al., 2022; Provenzi et al., 2018).

Finally, the results of the present study suggest that binaural spatialization of mother-voice and instruments triggered a relative event-related desynchronization compared to their monaural counterparts in theta, alpha and beta-band. Binaural speech and sound perception is age-dependent; while it optimizes in early adulthood, it remains a complex task for children (Nábělek & Robinson, 1982), and infants (Nozza et al., 1988). However, our results are in accordance with previous studies in newborn, showing the presence of neural correlates of spatial perception of auditory cues in the newborn brain (Németh et al., 2015). Our results are also reminiscent of the alpha-beta ERD observed in adults during passive listening of naturalistic spatialized auditory cues, suggesting that the oscillatory correlates spatialized naturalistic sound is already present in the neonatal period (Langiulli et al., 2023). The ERD observed might also reflect spatial attentional orienting toward the stimulus presented, as a frontal-central beta band ERD has been observed in adults during attentional focus toward spatially distinct speakers (Geirnaert et al., 2020).

Importantly, we observed that maternal voices presented in the binaural setting continued to elicit distinct alpha event-related synchronization compared to instrumental sounds. This observation suggests that the ERD observed in the binaural setting might reflect spatial auditory processing and attentional orienting processes occurring in parallel and distinct from mother-singing voice specific processing, with the former not interfering or competing with the latter in terms of oscillatory patterns.

To conclude and highlight the clinical relevance of this study, the broad-synchronization effect observed in response to the mother’s singing voice may have functional significance in the atypical neurodevelopmental processes associated with prematurity, thereby supporting early personalized interventions that encourage parents to sing to their hospitalized preterm newborns. The generation of synchronized neuronal activity has been linked to activity dependent self-organisation of developing networks, interneurons network maturation, as well as cortical and cortical-subcortical network functional connectivity (Ben-Ari, 2001; Duan et al., 2020; Khazipov & Luhmann, 2006; Khazipov & Milh, 2018). Brain wave synchronization is believed to facilitate functional connectivity and information processing, making it a key indicator of cortical network maturity (Gray et al., 1989). Preterm children present significant alteration of fMRI measured connectomes as well as thalamocortical connectivity compared to their full-term counterparts (Damaraju et al., 2010; Toulmin et al., 2021), differences that persist during adolescence and adulthood (Lahti et al., 2023; Papini et al., 2016). These differences are associated with motor and emotional deficits later during development.

In terms of EEG oscillatory activity, resting-state increased theta band connectivity and lower theta-alpha and beta power was observed in preterm compared to full-term infants, and this decreased power was correlated with behavioral outcomes (Kozhemiako et al., 2019). This low-frequency decrease in synchrony compared to full-term infants was also observed during memory and visual tasks (Moiseev et al., 2015). Oscillatory abnormalities in preterm infants persist during adolescence, with very preterm adolescent displaying decreased beta activity, and being more likely to present decreased power in theta-alpha-beta power in correlation with academic and motor outcomes (Twilhaar et al., 2019). Crucially, low-frequency oscillators in these frequency bands are precisely recruited in the current study by the mother-singing voice stimuli. Interestingly, functional connectome developmental trajectory in preterm infants has been shown to be improved by musical enrichment interventions, highlighting the clinical potential of investigating the underlying mechanisms of enrichment stimuli perception at a very young age in this vulnerable population (De Almeida et al., 2020; Lordier et al., 2019)(Van Der Veek et al., 2024).

Acting on the oscillatory pattern of these infants, using the knowledge derived by the previous study might then prove beneficial to devise interventions aimed at improving the clinical outcomes of very preterm infants.

## Limitations

An important limitation of the present study is the reduced sample size, which limits the generalizability of the findings to the broader preterm population. Moreover, the relatively small size of the premature infant’s scalp poses challenges in confidently isolating the activation of specific source. Additionally, including a sample of term newborns and an adult population could further validate the results and elucidate developmental differences potentially attributable to premature birth. Future analyses will involve typically term newborns and adults listening to the same sound stimuli, allowing for a comparison of neural responses across developmental stages and providing deeper insights into the influence of prematurity on auditory processing.

## Conclusions

In the present study, we showed that maternal singing represented a salient and distinctive auditory stimulus for preterm newborns. It synchronized the preterm infant’s brain, and elicited distinct neural responses compared to a stranger’s voice or an instrumental rendition of the same melody. In contrast, when presented in a binaural condition all auditory stimuli tend to elicit desynchronization responses in the brain. This supports the understanding that preterm infants exhibit a robust sensitivity to maternal singing voice, highlighting its potential in shaping early auditory, cognitive and affective development as an early protective intervention in a critical period of brain development.

## Acknowledgments

DB, JM and MF drafted the manuscript. MF and PSH supervised the project, revised and approved the final version. DB conducted the analyses. FBM and JM were responsible for participant recruitment and contributed to manuscript revisions. DG and LL provided critical revisions. AV composed the musical stimuli. SH was in charge of audio recording and the creation of auditory stimuli. All authors revised and approved the final version of the manuscript.

## Data availability statement

the datasets generated during the current study are available from the corresponding author on request.

## Ethics approval statement

The Swiss Research Ethics Committee approved the study (BASEC no. 2023-00718),

