## Supplementary material for "Maternal singing synchronizes the preterm infants’ brain": SI

***Additional Stimuli:***

Nine conditions were used in the experimental setup but only seven were discussed in the present paper. The two remaining conditions are described here: Gamma isochronic, i.e. background with superimposed isochronic gamma tones (tones at 73 Hz presented in each ear) and Gamma binaural, i.e. background with gamma tones presented at different frequencies in each ears.

***Faded Mother stimuli***

Faded binaural mother-voice stimuli was designed to test whether neonates could process their mother’s voice regardless of dB presentation. In these stimuli, the mother’s voice was faded into the vocal harmonic background. Significant contrasts between the faded mother voice and salient binaural mother voice are represented in SI Fig 5. Frontal cluster has been selected for representation in SI-Fig 5 for comparison with the Mother Binaural contrast.

Faded-binaural mother pattern of oscillatory activation was highly reminiscent of what was observed for the binaural mother voice. Indeed, as observed for binaural mother voice stimuli, faded mother voice elicited relative to background a relative ERD in the theta (>0.9s PSO, Frontal-Temporal), alpha (>1.9s PSO, RT-Frontal), and beta 20Hz bands (>0.9s PSO, Frontal, central, right -temporal) and broadband-gamma ERD (>0.9s, presenting its earliest and maximal amplitude effect in the temporal clusters, SI-Fig6). The only low-frequency exception is the occipital cluster displaying a delta-theta ERS post-pitch change. Interestingly, only weak and transient effects were observed in the delta-theta and beta bands when contrasting mother-binaural with faded mother stimuli (SI Fig 6), while a relative ERS is observed in the theta-alpha (>1.9s PSO, whole scalp for alpha, central-temporal for theta) and beta12 bands(>0.9s PSO, Frontal-RT-Occipital) when contrasting monaural mother’s-voice with faded mother. Interestingly, a mainly temporal broadband gamma ERD (over the whole HF spectrum) was observed for faded-mother compared to mother voice either monaural or binaural (SI-Fig6).

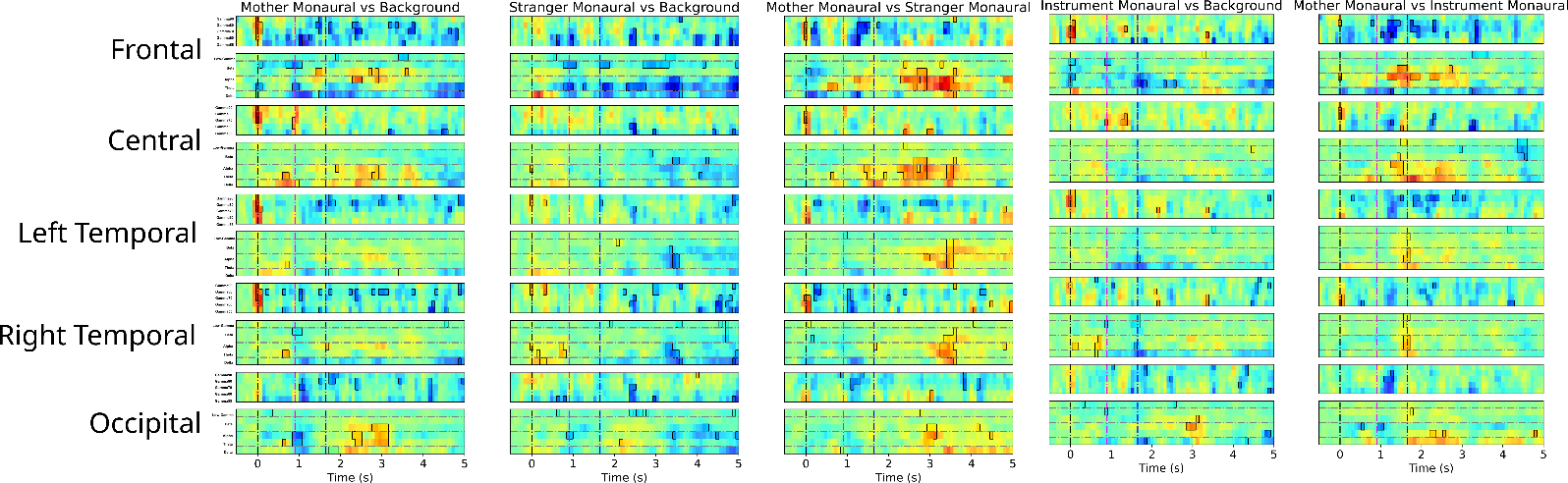

**SI Figure 1: Preterm infants’ EEG responses to monaural mother voice , stranger voice singing and instrument.** Contrasts between conditions are represented in the 5 columns. A simplified representation is presented with the average power for each frequency band statistically investigated. Framed areas represent time and frequencies where three statistical criteria are satisfied: First, absolute baseline corrected power exceeds 3 times the standard deviation of mean baseline power; Second, GLM analysis yielded a significant effect of Condition for this frequency band at this timepoint and Third, contrast analysis yielded a significant effect of the pairwise contrast considered after FDR correction. FDR: False Discovery Rate, GLM: General Linear Mixed Model.

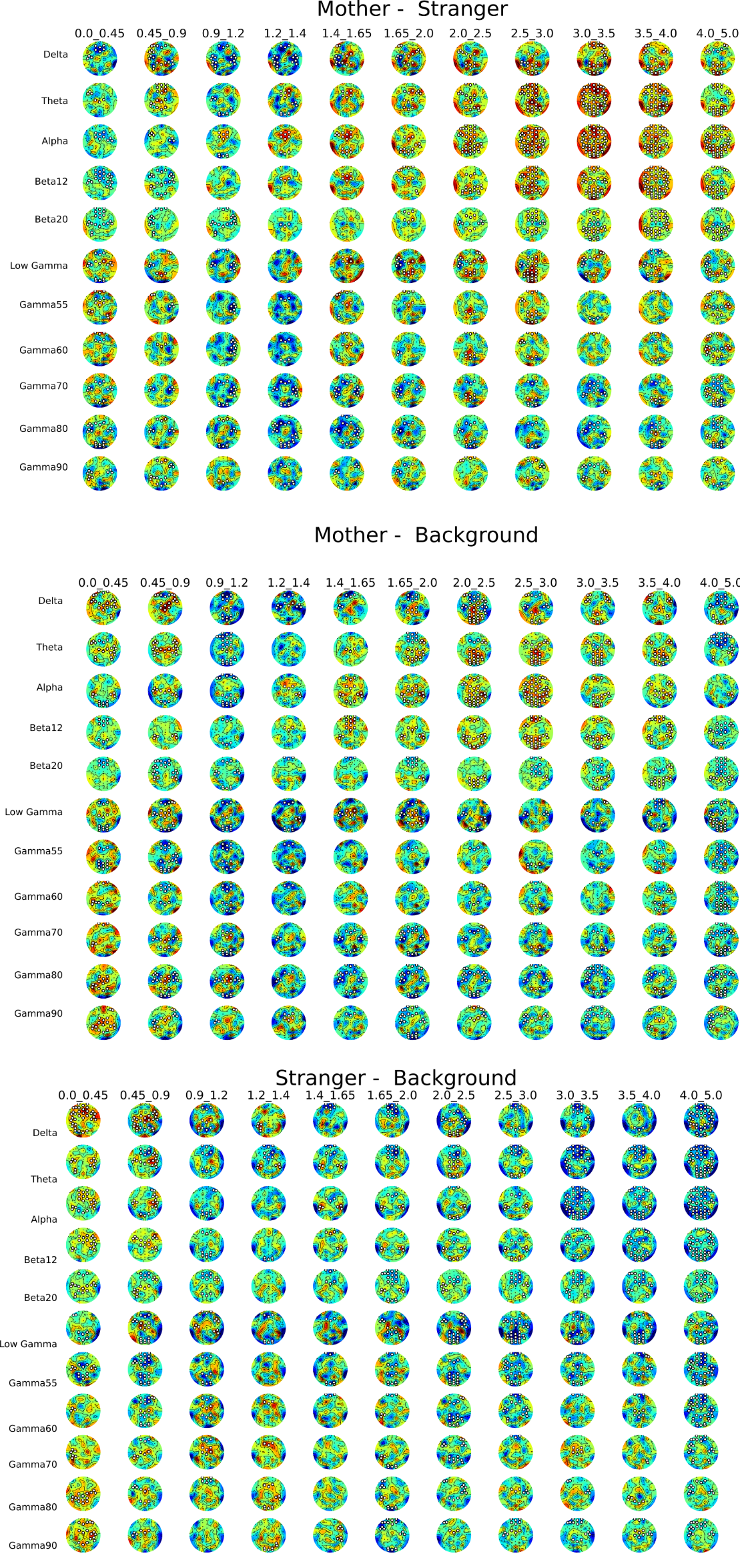

**SI Figure 2: Scalp topography of the monaural contrasts effect in premature infant’s brain.** White dots represent channels in a 3-channel micro-cluster where GLM analysis yielded a significant effect of Condition for this frequency band at this timepoint and contrast analysis yielded a significant effect of the pairwise contrast considered after FDR correction, thus providing a more topographically accurate location of significant effects. FDR: False Discovery Rate, GLM: General Linear Mixed.

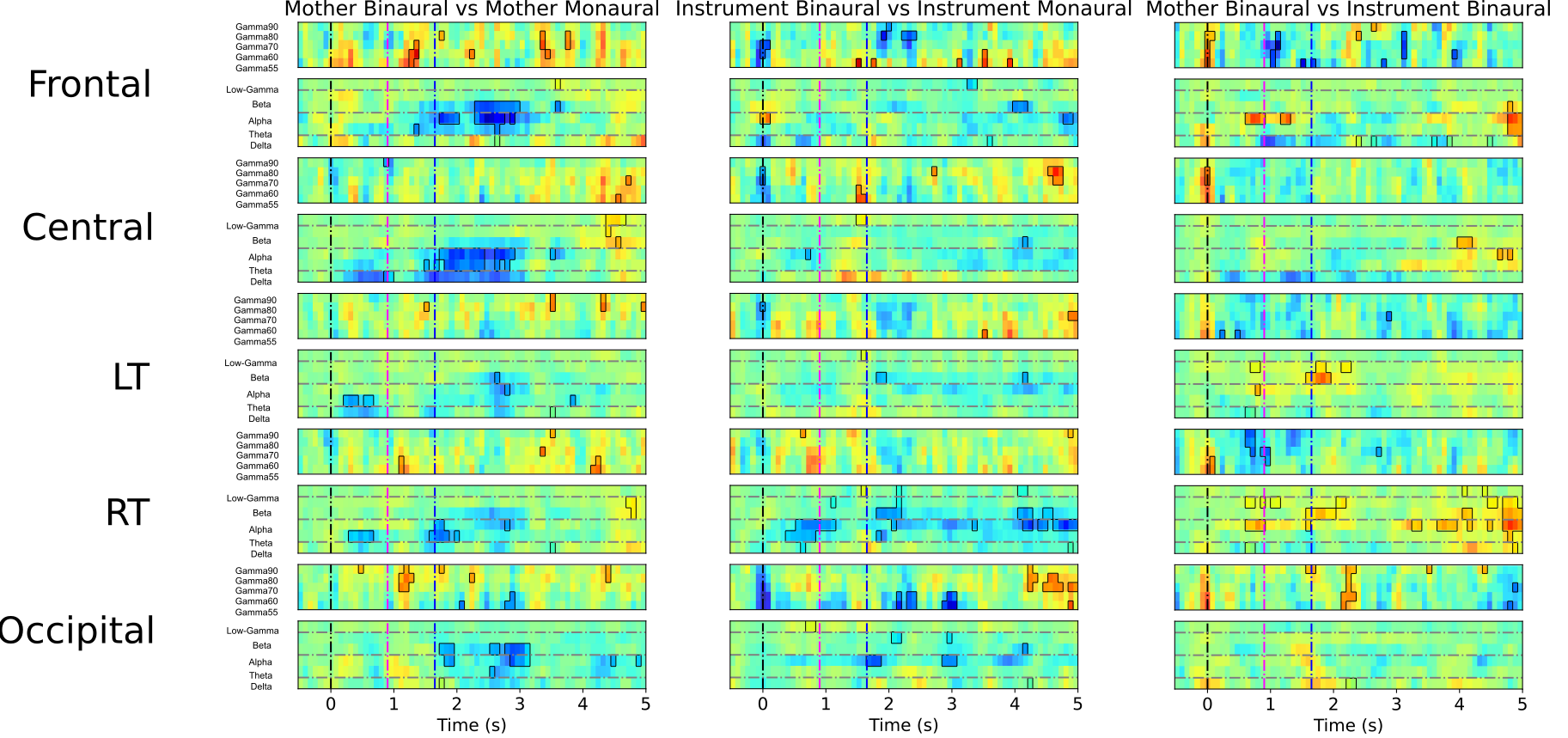

**SI Figure 3: Preterm infant’s EEG response to binaural mother voice and instrument.** Contrasts between conditions are represented in the 3 columns below. A simplified representation is presented with the average power for each frequency band statistically investigated. Circled areas represent time and frequencies where three statistical criteria are satisfied: First, absolute baseline corrected power exceeds 3 times the standard deviation of mean baseline power; Second,GLM analysis yielded a significant effect of Condition for this frequency band at this timepoint and third contrast analysis yielded a significant effect of the pairwise contrast considered after FDR correction

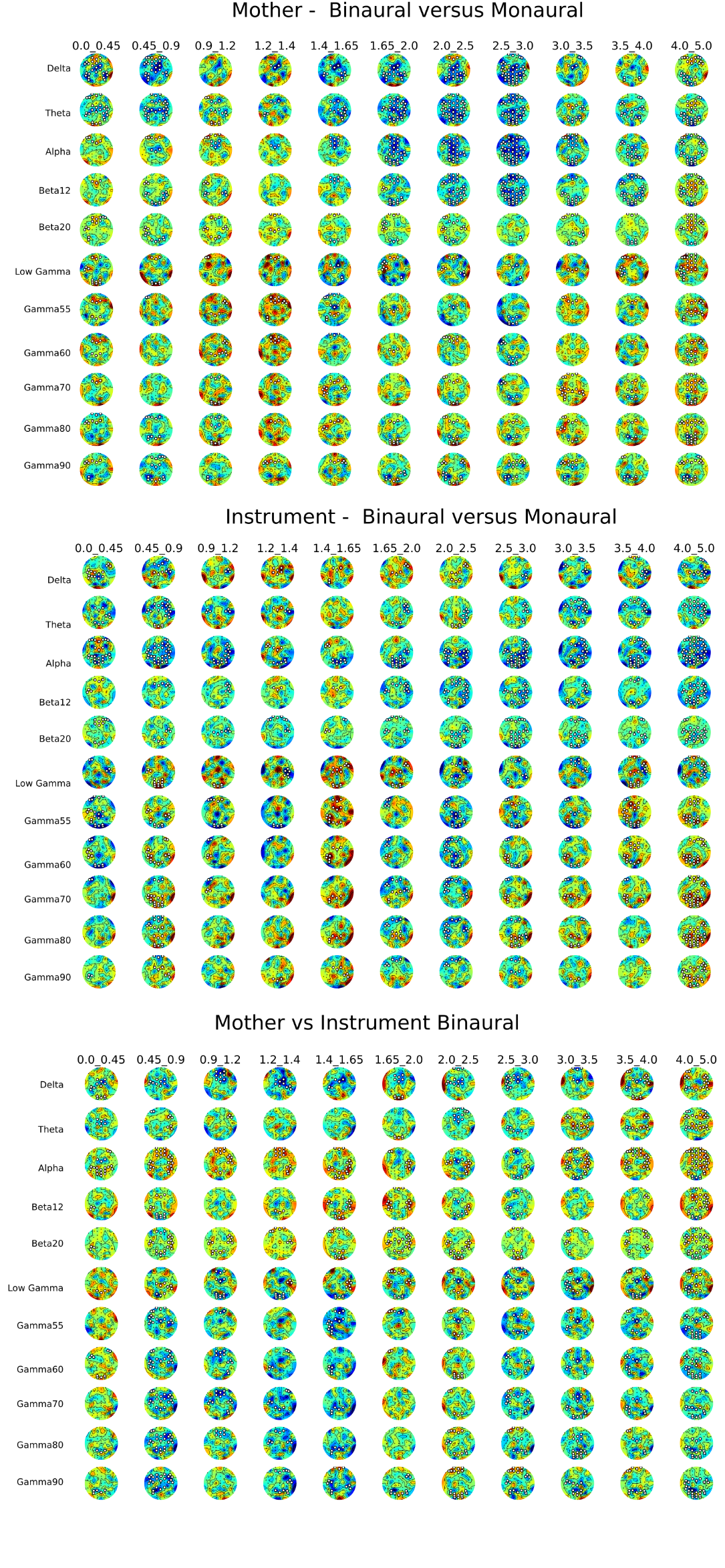

**SI Figure 4: Scalp topography of the binaural contrasts effect in preterm infant’s brain.** White dots represent channels in a 3-channel micro-cluster where GLM analysis yielded a significant effect of Condition for this frequency band at this timepoint and contrast analysis yielded a significant effect of the pairwise contrast considered after FDR correction, thus providing a more topographically accurate location of significant effects.

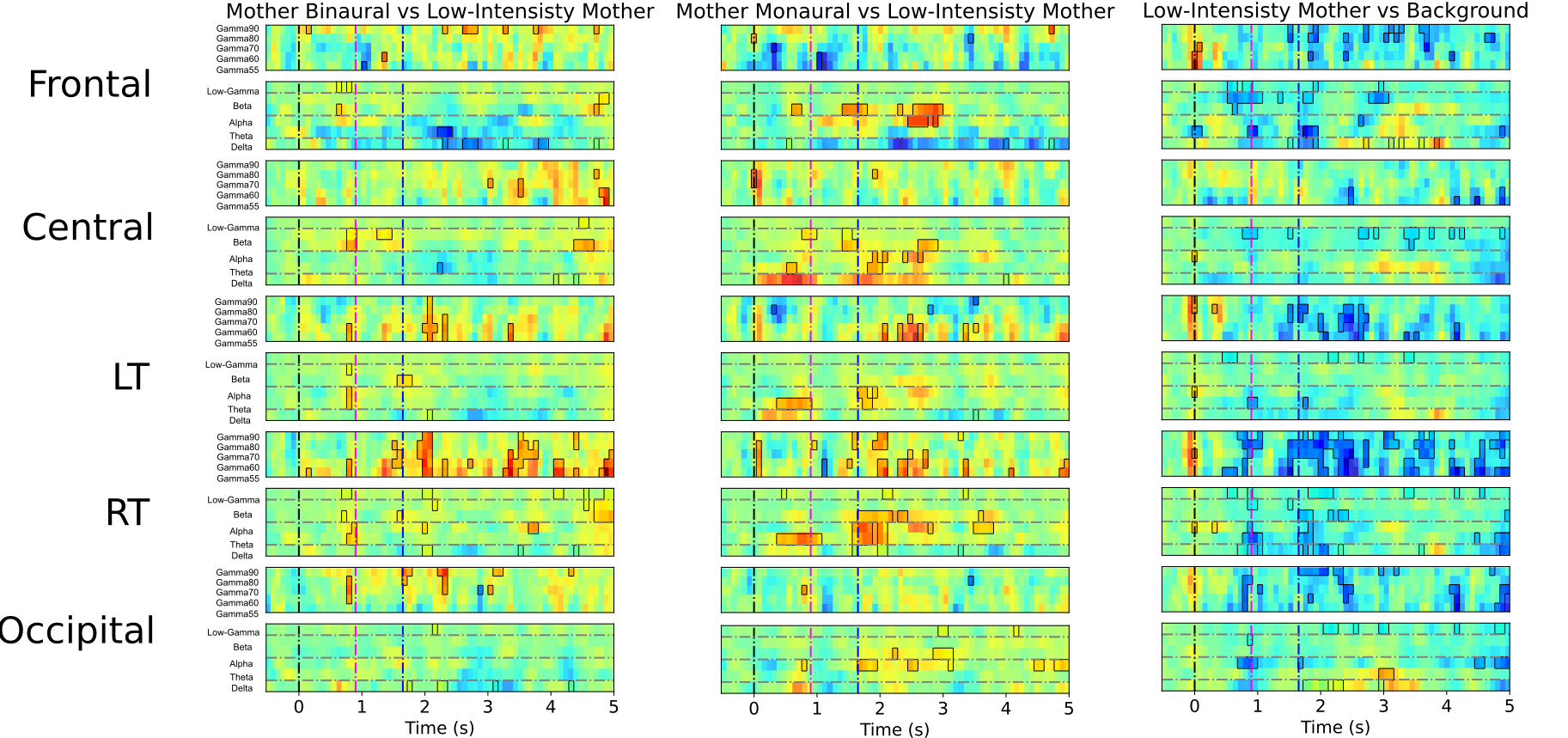

**SI Figure 5: EEG preterm infant’s response to faded-mother.** Contrasts between conditions for all ROIs are represented. A simplified representation is presented with the average power for each frequency band statistically investigated. Circled areas represent time and frequencies where three statistical criteria are satisfied: 1. absolute baseline corrected power exceeds 3 times the standard deviation of mean baseline power; 2. GLM analysis yielded a significant effect of Condition for this frequency band at this timepoint and 3. contrast analysis yielded a significant effect of the pairwise contrast considered after FDR correction.
